# Assessing the impacts of climate change on reproductive phenology in tropical rainforests of Southeast Asia

**DOI:** 10.1101/2021.08.24.457576

**Authors:** Shinya Numata, Koharu Yamaguchi, Masaaki Shimizu, Gen Sakurai, Ayaka Morimoto, Noraliza Alias, Nashatul Zaimah Noor Azman, Tetsuro Hosaka, Akiko Satake

**Affiliations:** Department of Tourism Science, Tokyo Metropolitan University, Tokyo 192-0397, Japan; Graduate School of Systems Life Science, Kyushu University, Fukuoka 819-0395, Japan; National Institute for Agro-Environmental Sciences, NARO, Tsukuba 305-8604, Japan; Forest Research Institute Malaysia, 52109 Kepong, Selangor, Malaysia; Graduate School of Advanced Science and Engineering, Hiroshima University, Hiroshima 739-8529, Japan; Department of Biology, Faculty of Science, Kyushu University, Fukuoka 819-0395, Japan

## Abstract

In humid forests in Southeast Asia, many species from dozens of plant families flower gregariously and fruit synchronously at irregular multi-year intervals^1–4^. Little is known about how climate change will impact these community-wide mass reproductive events. Here, we perform a comprehensive analysis of reproductive phenology and its environmental drivers based on a monthly reproductive phenology record from 210 species in 41 families in peninsular Malaysia. We find that the proportion of flowering and fruiting species decreased from 1976 to 2010. Using a phenology model with inputs obtained from general circulation models, we show that low-temperature flowering cues became less available during the monitoring period and will further decrease in the future, leading to decreased flowering opportunities in 57% of species in the Dipterocarpaceae family. Our results highlight the vulnerability of and variability in phenological responses across species in tropical ecosystems that differ from temperate and boreal biomes.

## Main

Simultaneous flowering in aseasonal forests in Southeast Asia, called general flowering, is one of the most spectacular and mysterious events that occur in tropical ecosystems. At irregular intervals of several years, diverse species, including species in the Dipterocarpaceae family, flower heavily^1–4^. Synchronous flowering and fruiting sometimes occur on a wide geographic scale (c. 10–10^9^ km^2^)^5–8^, together with flowering episodes occurring at other times at relatively small spatial scales^2,9^. A number of proximate cues for general flowering have been proposed, including drought^10–13^, the El Niño Southern Oscillation (ENSO)^6,11,14^, cloud-free conditions and high solar radiation^15–17^, and a night-time drop in the minimum temperature^2,5^. In addition, stored nutrients have been implicated as an endogenous factor regulating flowering^9,18,19^. Because recovery from nutrient shortages after heavy fruiting occurs relatively quickly within 1–2 years^20^, general flowering with intervals longer than 2 years is likely to be caused by external environmental factors rather than endogenous factors. Recent studies have demonstrated that the synergism between cool temperatures and drought is the major trigger for floral induction in dipterocarp trees^20–22^.

Global climate change brings elevated temperatures and more variable rainfall to Southeast Asia^23^. Projecting future phenological changes is an urgent task for the management and conservation of Southeast Asian rainforests because general flowering plays an important role in their successful regeneration and restoration^24,25^. However, the impact of climate change on the reproductive phenology of tropical rainforests is poorly understood^26–28^. This is partly due to the lack of long-term phenological data and the absence of predictive models that capture the mechanistic relationships between climatic factors and reproductive phenology in the tropics^26–28^.

To assess the past and future tropical phenology in Southeast Asia, we analysed historical records of reproductive phenology and meteorological data over 35 years in peninsular Malaysia and predicted what will happen regarding community-wide flowering events in the future. Our reproductive phenology data were collected from “Bulletin Fenologi Biji Benih dan Anak Benih (Bulletin of Seed and Seedling Phenology),” which was deposited at the library of the Forest Research Institute Malaysia (FRIM) located approximately 12 km northwest of Kuala Lumpur, Malaysia. Phenology monitoring was conducted based on monthly observations of the presence of flowers and fruits of tree species growing in the arboretum of FRIM from April 1976 to September 2010. After excluding species that did not satisfy the five required criteria for data accuracy, our phenology data included 210 species from 41 families (Extended Data 1). Dipterocarpaceae was the most abundant family (45% of total species), followed by Malvaceae (7.6%) and Leguminosae (6.2%) (Fig. 1a). These long-term phenology data from more than 200 species exposed to the same environment at the arboretum provide an excellent opportunity to compare phenological responses to climate change among species.

**Fig. 1.**
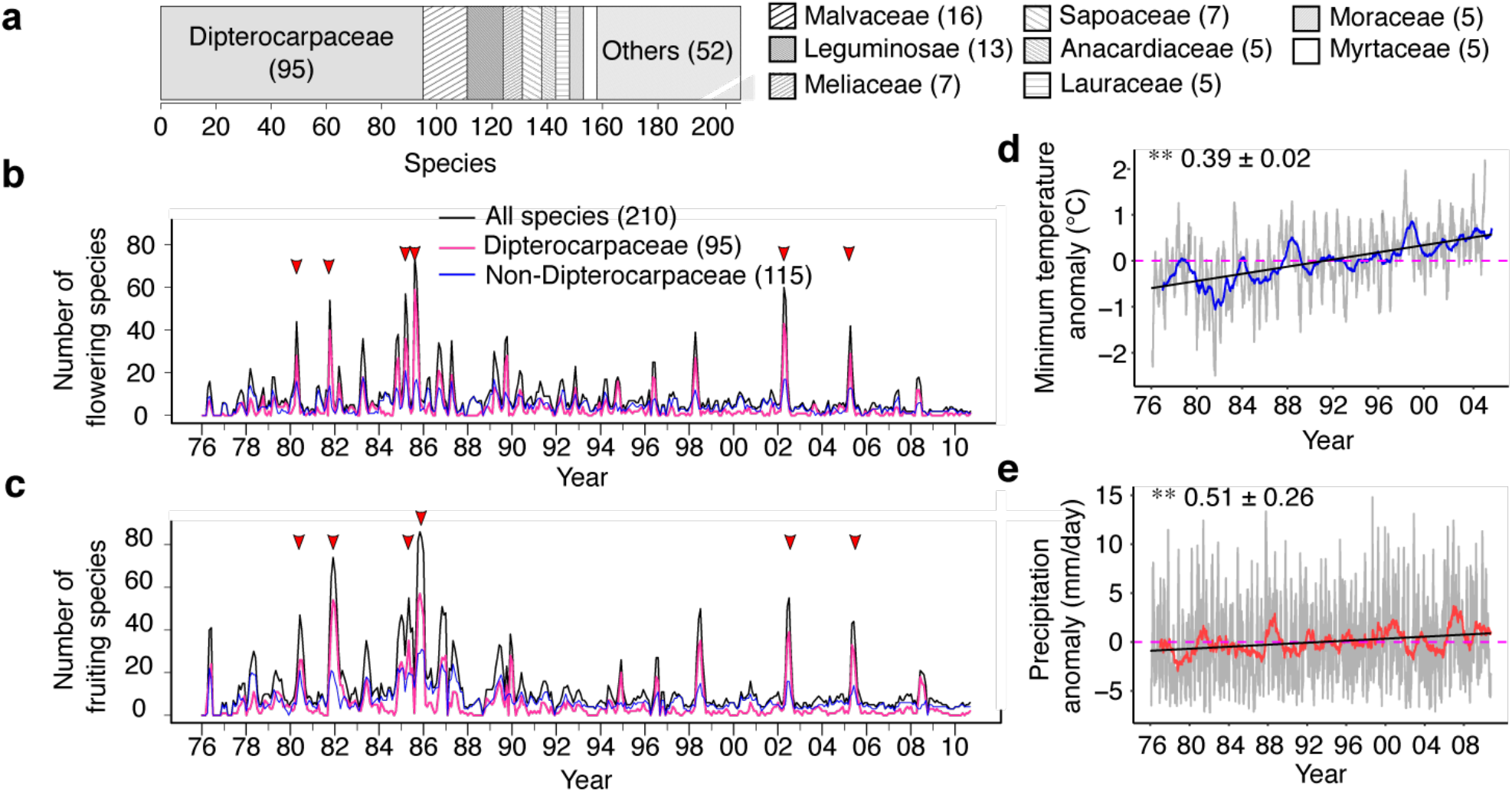
Flowering and fruiting records for 210 tropical tree species. **a**, Composition of species included in the phenology monitoring. The numbers in brackets indicate the number of species. The numbers of flowering (**b**) and fruiting (**c**) species among all species (black), dipterocarp species (pink), and non-dipterocarp species (blue) are shown. The horizontal arrows indicate mass flowering events, with more than 20% of monitored species blooming as flowers or fruits. Note that the number of monitored species was not always 210 due to missing values. **d**, A plot of the minimum temperature anomalies smoothed with the 30-day (grey) and 12-month (blue) moving averages. **e**, A plot of precipitation anomalies smoothed with the 30-day (grey) and 12-month (blue) moving averages. The black lines in panels **d** and **e** show the results of the linear regression of daily climatic variables, whereas the dashed pink lines indicate the normalized mean of each climatic variable over a monitoring period (1976–2005 for the minimum temperature and 1976–2008 for precipitation). The symbols indicate the results of the Mann– Kendall test. \*\**P* < 0.01. The mean change rate ± 95% of the confidence interval per decade is given for each plot.

The fractions of flowering and fruiting species fluctuated heavily between years. The greatest numbers of flowering and fruiting events occurred in 1985, in which more than 35% of monitored species participated in flowering and fruiting (Fig. 1b, c). Large flowering events with the flowering of more than 20% of monitored species occurred six times over 35 studied years, and these flowering events were followed by mass fruiting events (Fig. 1b, c, Extended Data Fig. 1a). These six large reproductive events at FRIM were synchronized with general flowering events monitored in natural forests in Peninsular Malaysia^8,9,23,24^, and the comparison suggests that the flowering and fruiting patterns between the arboretum at FRIM and natural forests were similar. The levels of between-species synchrony in flowering and fruiting events were significantly higher in Dipterocarpaceae species than in non-Dipterocarpaceae species (*P* < 0.0001, two-way analysis of variance (ANOVA), *n* = 4,465–6,555; Extended Data Fig. 1b). Moreover, the coefficients of variation in the proportion of flowering and fruiting species in Dipterocarpaceae were twice as large as those in non-Dipterocarpaceae species (1.787 and 1.583 for Dipterocarpaceae and 0.803 and 0.753 for non-Dipterocarpaceae, respectively). These results indicate that Dipterocarpaceae plays a pivotal role in general flowering.

We detected decreasing trends in the proportions of flowering (*P* = 0.0021, Mann-Kendall (MK) test, two-sided, *n* = 400; Extended Data Table 1) and fruiting species (*P* < 0.0001, MK test, *n* = 400; Extended Data Table 1) from the mid-1970s to the early 2000s. In contrast, temperature and precipitation during this period revealed an increasing trend at mean rates of 0.39 ± 0.02°C (*P* < 0.0001, MK test, two-sided, *n* = 10,330; Fig. 2d) and 0.51 ± 0.26 mm/day per decade (*P* = 0.027, MK test, two-sided, *n* = 12,668; Fig. 2e), respectively. The monthly frequency of flowering and fruiting varied largely among species (Fig. 2a, b); 17% of species flowered at least once per year, whereas 25% of species flowered only once every 10 years (Extended Data 2). Regardless of this large variation in the frequency of reproductive events across species, most species exhibited clear reproductive seasonality. At the community level, two flowering peaks occurred in April and October (Fig. 2a), followed by fruiting peaks in June and December, respectively (Fig. 2b). The seed dispersal time, which was calculated as the month when all mature fruits dropped from their mother tree, peaked in February or August, two months after the fruiting peaks (Fig. 2c). The timing of the two seed dispersal peaks matched the phases in which temperatures and rainfall started to increase (Fig. 2d, e), which was consistent with the finding that seed dispersal is timed to match the favourable humidity condition for the survival of offspring^29^. Among the nine families containing at least five species, only Moraceae, which includes the genus *Ficus*, produced flowers and fruits almost year-round without any seasonality (Fig. 3). Other families show synchronized flowering phenology in both spring and autumn (e.g., Dipterocarpaceae; Fig. 3) or spring flowering dominance (e.g., Anacardiaceae, Lauraceae, and Meliaceae; Fig. 3).

**Fig. 2.**
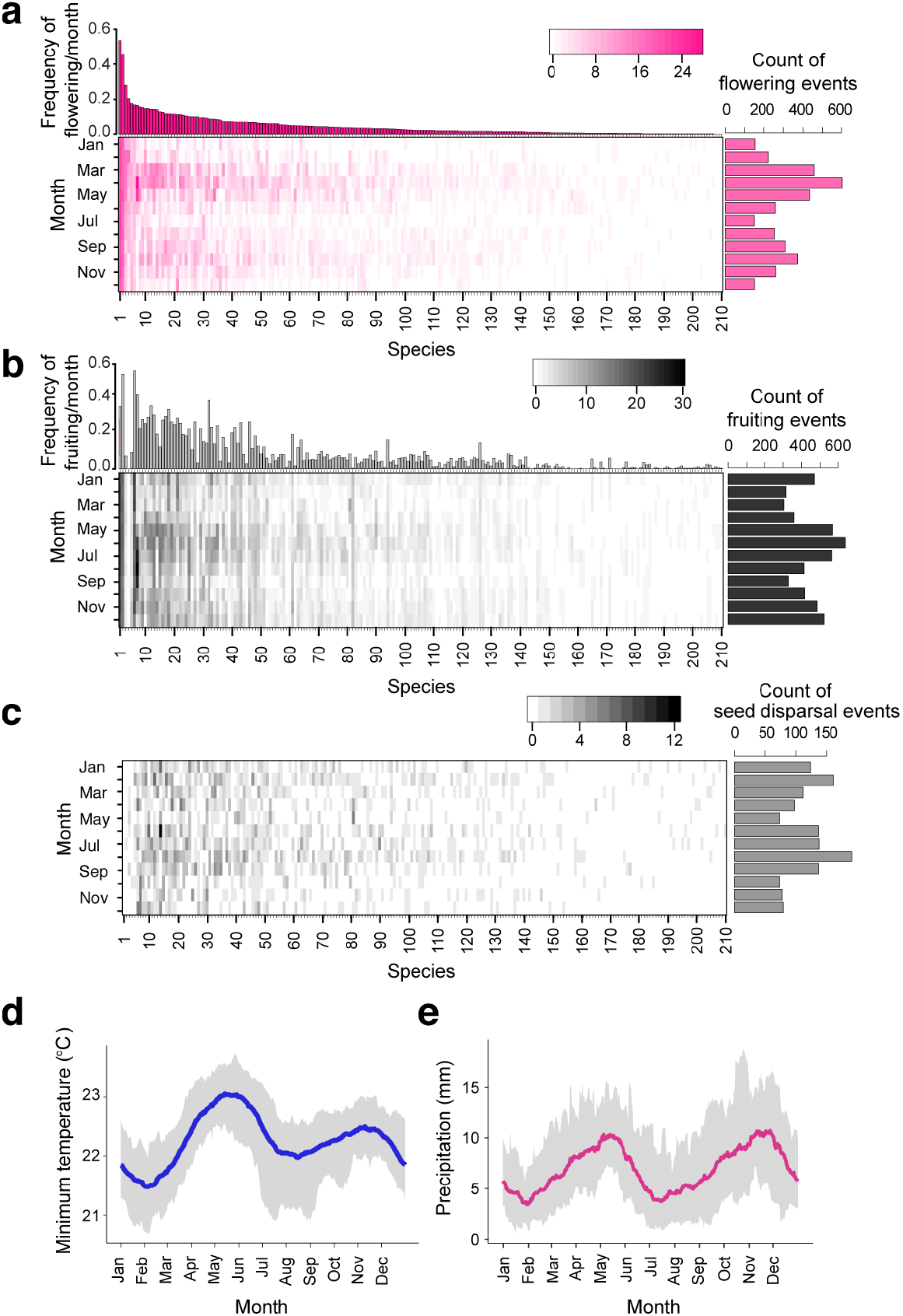
Frequency and seasonality of reproductive events. The frequency and seasonal trends of flowering (**a**), fruiting (**b**), and seed dispersal (**c**) events in 210 tropical tree species. The number given to each species corresponds to the species name shown in Extended Data 2. **d**, Average (line) ± standard error (envelope) of temperature calculated from the 30-day running means during 1976–2010. **e**, Average (line) ± standard error (envelope) of precipitation calculated from the 30-day running means during 1976–2010.

**Fig. 3.**
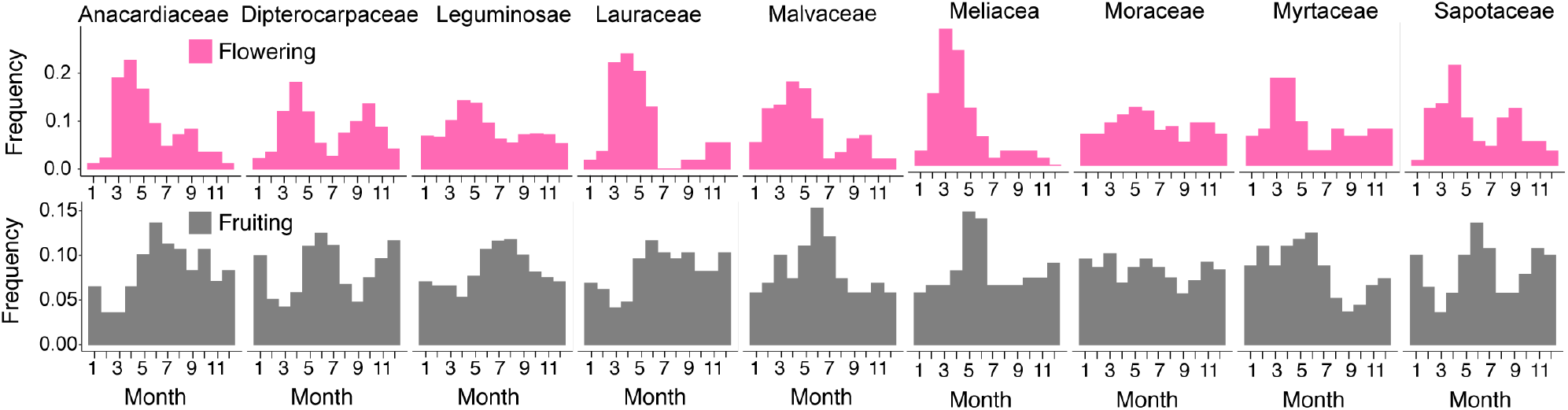
Seasonal trends in flowering and fruiting phenology. Histograms of flowering (pink) and fruiting (grey) frequencies were drawn for nine families in which at least five species were included. The number of species included for each family is given in Fig. 1a.

The decreased proportions of flowering and fruiting species observed in the past can be explained by increased temperatures or decreased drought event frequencies because cool temperatures and drought have been suggested to be major environmental drivers of general flowering^20–22^. To investigate the relationships between flowering and temperature and between flowering and precipitation, we adopted a model that was developed to predict flowering phenology in Dipterocarpaceae^21^ and was further extended to predict the flowering phenology of tropical plants on Barro Colorado Island, Panama^30^. The previous model was successful in explaining the weekly flowering phenology of five dipterocarp species over 10 years in the Pasoh forest reserve in peninsular Malaysia; thus, it is a reliable model for forecasting future flowering phenology in Dipterocarpaceae. The model assumed that potential environmental cues to floral induction, the cool unit (CU) and drought unit (DU), accumulate for *n*_1_ days prior to the onset of floral induction, and flowers then develop for *n*_2_ days before opening (Extended Data Fig. 2). The DU model evaluates whether drought alone cues flowering (drought-induced flowering). The CU × DU model evaluates whether cold and drought have a multiplicative effect on flowering (low temperature- and drought-induced flowering). Logistic regression was performed using only DU and using CU × DU as the explanatory variables and using the presence or absence of a first flowering event as the dependent variable. We examined these two models because dipterocarp flowering has been predicted successfully by both of these models in previous studies^21,29^.

To perform model fitting, we focused only on Dipterocarpaceae because of the low percentage of missing values (4.81%) compared to those of non-Dipterocarpaceae species (16.8% on average; Extended Data 1). Because some species have similar flowering phenology, we first performed time-series clustering using 98 dipterocarp species based on the similarity of their flowering phenology (Fig. 4a) and then carried out forward selection of the optimal number of phenological clusters based on the minimization of the Akaike information criterion (AIC) (Extended Data Fig. 3). We used data from May 1976 to March 1996 to train the model and data from July 1997 to April 2005 to validate the model. We chose these periods for the model training and validation because there was a blank period in the dataset from April 1996 to June 1997 in which climate data were missing.

**Fig. 4.**
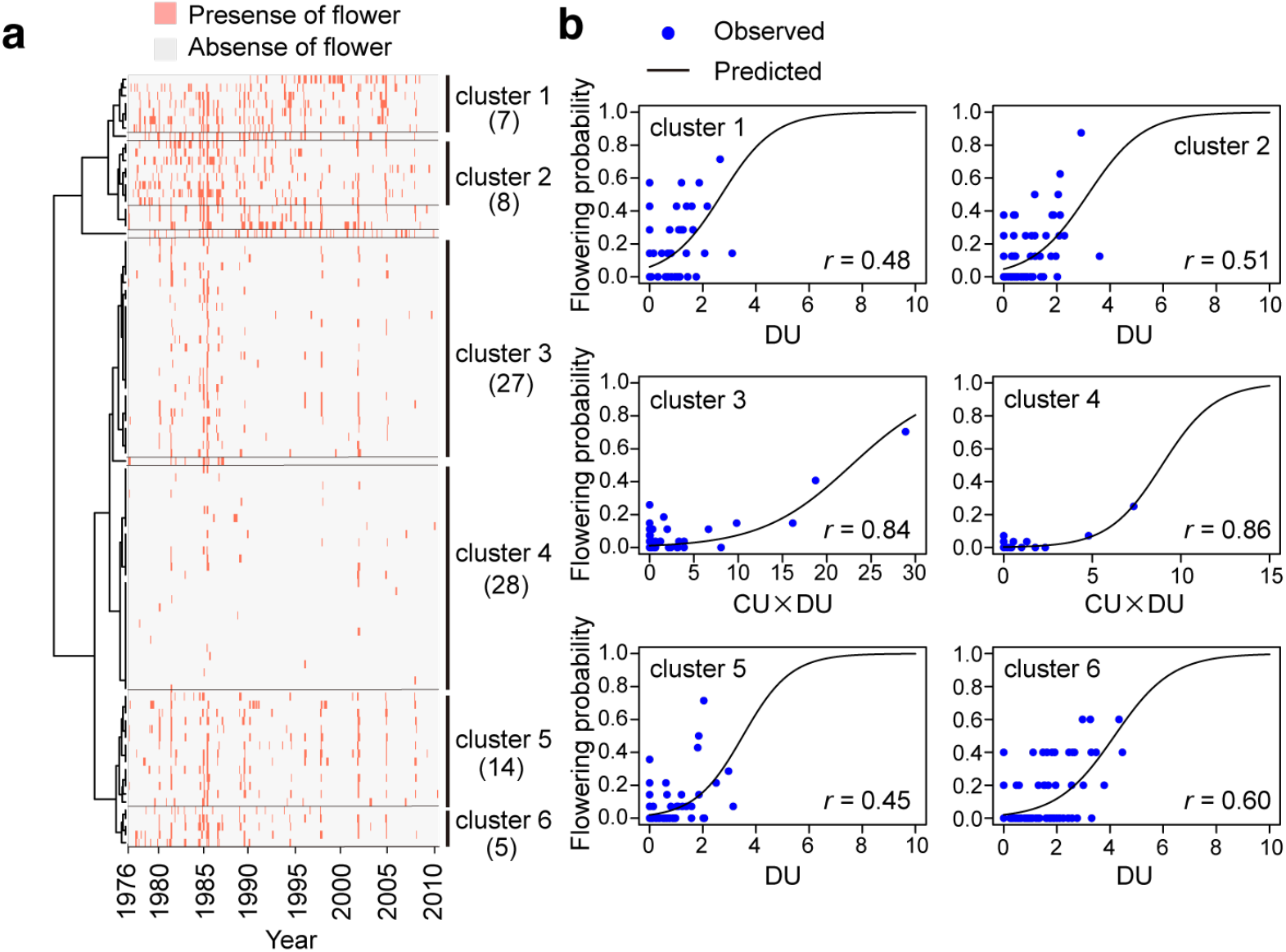
Clustering of phenological patterns and flowering phenology prediction for each phenological cluster. **a**, Time-series clustering of 95 dipterocarp species based on the similarity of their flowering (pink) and nonflowering (grey) patterns. The optimal number of clusters 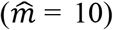 was identified by minimization of AIC. Among the 10 clusters, clusters with fewer than four species were excluded for phenology forecasting because of their small sample sizes. Thus, six clusters were used for phenology forecasting. The numbers in brackets indicate the number of species included in each cluster. **b**, Comparisons between the predicted (black) and observed (blue) flowering phenology of each phenological cluster during May 1976–March 1996. To explain flowering phenology, the CU × DU model was applied for clusters 3 and 4, while the DU model was selected for the other clusters. Pearson’s correlation coefficient (*r*) is given for each plot. Details for the model fitting results are given in Extended Data Table 3.

The model fitting results revealed that the optimal number of phenological clusters was 10 (seven clusters and three independent species; Extended Data 3). After removing independent species and clusters with fewer than 5 species (due to their small sample sizes), six clusters remained (Fig. 4a; Extended Data 4). The CU × DU model was selected to explain the flowering phenology of clusters 3 and 4, two major clusters including 27 and 28 species, respectively (Fig. 4b; Extended Data Table 2). The flowering phenology of other clusters was explained by drought only (Fig. 4b; Extended Data Table 2). The area under the ROC (receiver operating characteristic) curve (AUC) values ranged from 0.64–0.78 for the training data and 0.62–0.79 for the validation data (Extended Data Table 2), suggesting an acceptable discrimination ability of the model for most clusters^31^. Because cluster 1 showed low AUC values (0.64 for the model training and 0.62 for the model validation), predictions of this cluster must be approached with caution (Extended Data Table 2).

We predicted the future flowering phenology based on the model for each of six phenological clusters under two climate scenarios, Representative Concentration Pathways (RCP) 2.6 with limited CO_2_ emissions and RCP 8.5 with high CO_2_ emissions, which were simulated using three general circulation models (GCMs; GFDL–ESM2M; Fig. 4a, b, IPSL–CM5A-LR and MIROC5; Extended Data Fig. 4). Compared to the 1976–2005 period, the minimum temperature was predicted to increase by 1.2 ± 1.1°C under the RCP2.6 scenario and by 3.1 ± 1.7°C under the RCP8.5 scenario by 2099. Under the RCP2.6 scenario, the mean predicted flowering probabilities during the 2050–2099 period across the three models decreased in clusters 3 and 4 to 57% and 49% of the predicted flowering probabilities during the 1976–1996 period, respectively (Fig. 5c). Under the RCP8.5 scenario, the mean predicted flowering probabilities in clusters 3 and 4 were further reduced to 37% and 28% during the 2050–2099 period, respectively (Fig. 5c). The decreased flowering probabilities in these two phenological clusters were caused by lower occurrences of low-temperature flowering cues in the future (Extended Data Fig. 5). Under the RCP8.5 scenario, low-temperature signals that trigger flowering in the species included in phenological clusters 3 and 4 rarely occurred or completely disappeared (Extended Data Fig. 5). In contrast, in species included in other clusters that were sensitive only to drought cues for flowering, the mean flowering probabilities were unchanged between the 1976–1996 and 2050– 2099 periods (Fig. 5c) because the drought-related flowering cues were rather stable throughout the simulations (Extended Data Fig. 5).

**Fig. 5.**
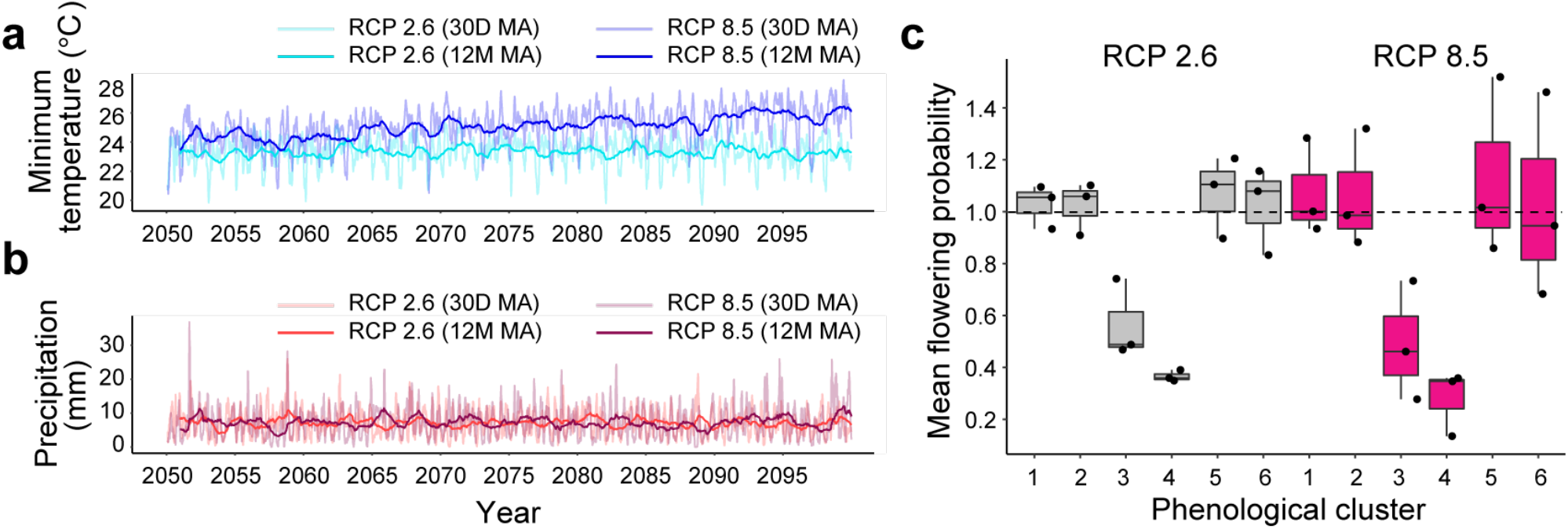
Prediction of future flowering phenology under two climate scenarios (RCP2.6 and RCP8.5) Minimum temperature (**a**) and precipitation (**b**) simulated by GDFL–EMS2M for 2050–2099. The 30-day (30 D) and 12-month (12 M) moving averages (MAs) were plotted. **(c)** Box plots of normalized flowering probabilities predicted for 2050–2099 for each phenological cluster under two climate scenarios. Predictions from three GCMs (GDFL–EMS2M, IPSL–CM5A-LR, and MIROC5) were plotted as points. For each model, the prediction was normalized using the 1976– 1996 phenology of each phenological cluster. A dotted line indicates the level that is equal to the one for 1976–1996 period. The horizontal line inside each box and the length of each box indicate the median and the interquartile range (the range between the 25th and 75th percentiles), respectively. The whiskers indicate points within 1.5 times the interquartile range.

To confirm our predictions, we extended our phenology forecasting to three other regions in Southeast Asia, Trang Province in Thailand^32^, Lambir Hills National Park in Malaysia^12^, and central Kalimantan in Indonesia^13^, where long-term phenology monitoring plots exist (Fig. 6a). Decreased flowering probabilities were predicted only in clusters 3 and 4 in all regions, while the flowering probabilities of other species were predicted to be robust (Extended Data Fig. 6), confirming the predictions obtained in FRIM. A comparison of seasonal flowering patterns along a latitudinal gradient based on historical climate data simulated during the 1976–1996 period from GCMs revealed shifts from a unimodal flowering peak in spring (March in the northern hemisphere) in Trang Province, bimodal or weak flowering peaks in spring and fall in FRIM and Lambir National Park and a pronounced flowering peak only in spring (September in the south hemisphere) in central Kalimantan (Fig. 6b). The predicted seasonal flowering patterns are consistent with previous observations^12,13,32^, showing that using low temperatures and drought to forecast phenology can be applicable to wide regions in Southeast Asia. The latitudinal gradient of seasonal flowering patterns was predicted to be robust to climate change in the 21st century (Fig. 6c), suggesting that the different seasonal distribution of flowering probability among the four regions can be explained by the differential seasonal rainfall patterns across the regions (Extended Data Fig. 7). These results suggest that the phenological responses of tropical trees in Southeast Asia to climate change are not qualitative but are rather quantitative.

**Fig. 6.**
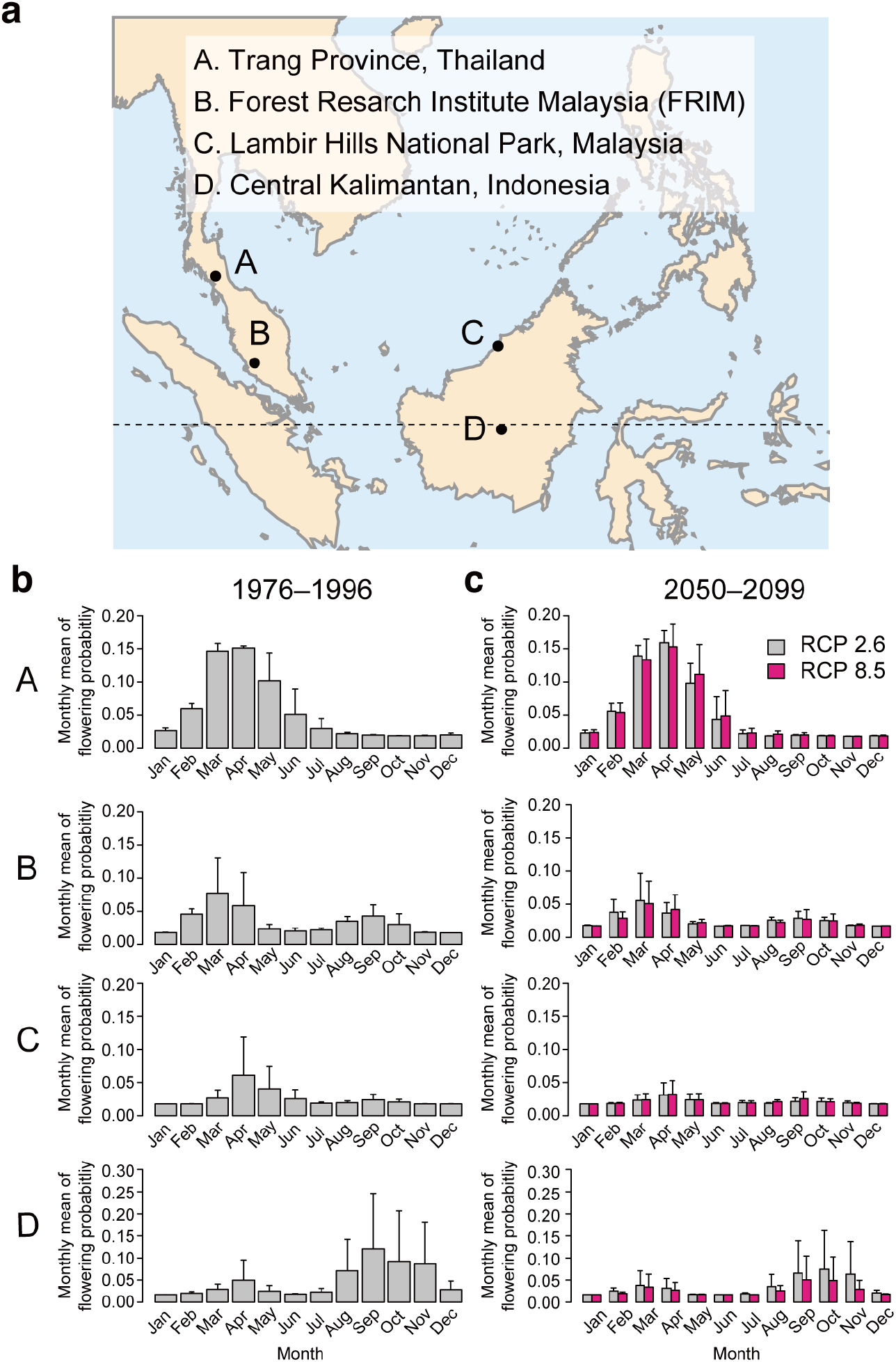
Comparison of predicted flowering phenology across four regions in Southeast Asia. **a**, A map illustrating the four study sites (A–D) used for phenology forecasting. The dotted line indicates the equator. **b**, Means ± standard errors of the monthly flowering probability predicted during the 1976–1996 period. **c**, Means ± standard errors of the monthly flowering probability predicted during the 2050–2099 period in the four studied regions. The means and standard errors across three GCMs were calculated from predictions by GDFL–EMS2M, IPSL–CM5A-LR, and MIROC5. A–D correspond to each of four study sites in the map (**a**).

Phenological shifts are among the most widely studied biotic responses to climate change^33^. Most phenological shift observations come from temperate and boreal biomes where advancing biological springs and delayed biological winter arrivals are documented in response to increased temperatures^34,35^. Tropical species have been suggested to be more sensitive to climatic changes than temperate and boreal species because they have evolved in areas with less seasonal environmental variation^36,37^. However, no studies have addressed the vulnerability of tropical species under future climate change. Our projections of 21st century changes in the flowering phenology of tropical plant species revealed that a 1.2°C increase in temperature under the RCP2.6 scenario resulted in an approximately 50% decrease in the future flowering probabilities of 57% of dipterocarp species that are sensitive to low-temperature flowering cues. In a temperate perennial herb that requires winter cold for floral initiation, a significantly decreased flowering probability was predicted when the temperature increased by 4.5°C^38^. These results suggest that tropical species are more sensitive and vulnerable to climate change than species located in temperate ecosystems. Our results also highlight the variable features of phenological changes among species in response to climate change. Forty-three percent of dipterocarp species are predicted to be sensitive only to drought for flowering, and their reproductive activities are robust to climate change. Different phenological responses across species would alter forest regeneration and, eventually, the plant species composition in the future. Regardless of the significant effect of climate change on the flowering probability at the quantitative level, the seasonal distribution of flowering probability were found to be robust in wide regions of Southeast Asia; this result represents another interesting difference between tropical and temperate plant species.

Although the data presented here are one of the longest records of reproductive phenology in tropical ecosystems^39^, we still need to be cautious when interpreting these results because there is room to extend our analyses from the phenological cluster level to the species level when longer-term and higher-temporal-resolution data become available. With upgraded phenological data, the estimation accuracy of the environmental drivers of tropical phenology and predictive power will be improved. Because observations of reproductive phenology in tropical plants are still rare due to a paucity of long-term studies^26–28^, continued phenology monitoring is necessary^39^.

There have been only a few reports on past phenological changes in tropical plants. In Kibale National Park, Uganda, fruiting phenology from two data sets (covering 1970–1983 and 1990–2002) revealed that the proportion of individual fruiting was negatively correlated with the minimum temperature and that increased rainfall was associated with the complete absence of fruiting in common tree species^40^. In Ranomafana National Park, Madagascar, 12 years of fruiting phenology observations showed a correlation between increased rainfall and an increase in fruiting in some tree species^41^. These studies highlight the complex and variable features of phenological changes among tropical plant species in response to climate change^37^. An improved mechanistic understanding of the environmental drivers of reproductive phenology in diverse species in different tropical ecosystems will unravel the complex nature of phenological responses in the tropics and will allow the extension of future reproductive phenology projections from regional to global scales^42^.

The rapid global warming that occurred over the last millennium was unprecedented^43^. Our results suggest that plastic responses to climate change at the individual level may not be high for the tropical species studied herein. Moreover, species with long generation times are the least likely to be rescued by evolution alone^44^. Early detections of biotic change signatures and predictions of the magnitude and direction of changes in plant reproductive phenology will benefit management programmes and aid in the sustainable future of the most diverse ecosystems on Earth.

## Methods

### Data collection of flowering and fruiting phenology

Monthly reproductive phenology data recorded over 35 years were collected from the “Bulletin Fenologi Biji Benih dan Anak Benih (Bulletin of Seed and Seedling Phenology),” which was deposited at the FRIM library. The bulletin reported seed and seedling availabilities and the flowering and fruiting phenology of trees at several research stations in Malaysia. The present study collected flowering and fruiting records of trees grown in FRIM arboretums located approximately 12 km northwest of Kuala Lumpur, Malaysia (latitude 3°24 ‘N, longitude 101°63 ‘E, elevation 80 m). There are both dipterocarp and non-dipterocarp arboretums in FRIM, both of which were founded in 1929. These arboretums preserve and maintain living trees for research and other purposes. Each month, three research staff members of FRIM with sufficient phenology monitoring training made observations with binoculars to record the presence of flowers and fruits on trees of each species on the forest floor from April 1976 to September 2010. The phenological status of the trees was recorded as “flowering” during the developmental stages from flower budding to blooming and as “fruiting” during the developmental stages from the occurrence of immature fruit to fruit ripening. Because only one or two individuals per species are grown at the FRIM arboretums, the flowering and fruiting phenology were monitored using these individuals. The resultant flowering and fruiting phenology data included a time series of binary data (1 for presence and 0 for absence) with a length of 417 months.

The original data included 112 dipterocarp and 240 non-dipterocarp species. We excluded 17 dipterocarp and 125 non-dipterocarp species based on the following five criteria for data accuracy.

1. Percentage of missing values is less than or equal to 50%: If the monthly flowering or fruiting phenology data of a species included a substantially large number of missing values (more than 50%), the species was excluded.
2. Stable flowering period: We considered an observation to be unreliable if the flowering period was significantly different among flowering events (if the coefficient of variation in the flowering period was larger or equal to 1.0).
3. Flowering period is shorter than or equal to 12 months: We considered an observation to be unreliable if the flowering period was longer than 12 months because it was unlikely that the same tree would flower continuously for longer than 1 year.
4. The flowering and fruiting frequencies were not significantly different between the first and second half of the census period: When the flowering frequency was zero for the first half of the observation period but was larger than 0.1 for the second half of the observation period, it was likely that the specimen was newly planted after monitoring began. Similarly, when the flowering frequency was zero for the second half of the observation period but was larger than 0.1 for the first half of the observation period, it was likely that the species died during monitoring. We removed these species for our analyses. We adopted the same criteria for the fruiting phenology data.
5. We removed overlapping species, herb species, and specimens with unknown species names.

After removing unreliable species based on the five criteria explained above, we obtained 95 dipterocarp and 115 non-dipterocarp species (Extended Data 1). We used these species for further analyses.

### Detection of seasonality in reproductive phenology

To compare the flowering and fruiting phenology seasonality among different families, nine families that included at least five species were used. The number of flowering or fruiting events was counted for each month from January to December during a census, and then the frequency distribution was drawn as a histogram. Similarly, we also generated a histogram for the seed dispersal month, which was calculated as the month when fruiting ended (i.e., when the binary fruiting phenology data changed from one to zero).

### Classification of phenological patterns

To classify the phenological patterns, we performed time-series clustering using the R package “TSclust”^48^ with the hierarchical clustering method based on the Dynamic Time Warping (DTW) distance of the flowering phenology data of each species. For this analysis, time points at which there were missing values for at least one species were excluded. Because of the large number of missing values in non-Dipterocarpaceae species, we performed time series clustering only for the Dipterocarpaceae species based on 394 time points in total. The number of phenological clusters was estimated based on AIC, as explained below.

### Climate data

Daily minimum, mean, and maximum temperatures and precipitation data monitored at the FRIM KEPONG (3° 14’ N, 101° 42’ E, elevation 97 m) weather station were provided by the Malaysian Meteorological Department. We used the daily minimum temperature for our analysis because there were fewer missing values compared to the numbers of missing daily mean and daily maximum temperature values. The periods in which climate data were available were from March 1, 1973 to March 31, 1996, and from July 23, 1997 to April 20, 2005. We removed periods in which there were missing values spanning longer than five days. When the range of missing values spanned a period shorter than 3 days, we approximated these missing values using the mean minimum temperatures recorded on the adjacent three days. Although solar radiation data were not available for our study, the use of precipitation is sufficient for model fitting because there is a significant negative correlation between solar radiation and precipitation in Southeast Asia^49^.

### Climate data generated by general circulation models (GCMs)

As the future climate inputs, we used bias-corrected climate input data from January 1, 2050, to December 31, 2099, with a daily temporal resolution and a 0.5° spatial resolution, provided by the ISI-MIP project^50^; these data are based on the Coupled Model Intercomparison Project Phase 5 (CMIP5) outputs from three GCMs: GFDL–ESM2M, IPSL–CM5A-LR, and MIROC5. To compare the flowering phenology between 1976–1996 and 2050–2099, bias-corrected GCM data from May 1, 1976, to March 31, 1996, were also used. This period (May 1, 1976–March 31, 1996) is consistent with the period used for model fitting. We selected daily minimum temperature and precipitation time series from the 0.5° grid cells corresponding to the study site for phenology monitoring at FRIM. To compare flowering phenology among regions, we also used the same set of data from three other regions in Southeast Asia: Trang Province in Thailand (7° 4’ N, 99° 47’ E), Lambir Hills National Park in Malaysia (4° 2’ N, 113° 50’ E), and central Kalimantan in Indonesia (0° 06’ S, 114° 0’ E). Because the study site in FRIM was not in the centre of a 0.5° grid cell, we interpolated the data using four grid cells in the vicinity of the observation site. We used the weighted average according to the distance between each observation site and the centre of each corresponding grid cell.

Although the climate input data provided by ISI-MIP were already bias-corrected, we conducted additional bias correction at FRIM using a historical scenario for each GCM dataset and the observed weather data from January 1, 1976 to December 31, 2004 based on previously presented protocol^51^. We did not implement any bias correction for the frequency of dry days or precipitation intensity of wet days^51^ because we only focused on the average precipitation.

The variances in the annual fluctuation of the monthly mean precipitation were not the same between the observation data and historical GCM runs at FRIM. For all three GCMs (GFDL– ESM2M, IPSL–CM5A-LR, and MIROC5), the variances in the yearly fluctuation output by the GCMs tended to be larger than that of the observed data at the FRIM KEPONG weather station during winter and spring. On the other hand, during summer and fall, the variances output by the GCMs tended to be smaller than that of the observed data. These biases could not be corrected using the previous method^51^. Therefore, we conducted the following bias correction for these data:

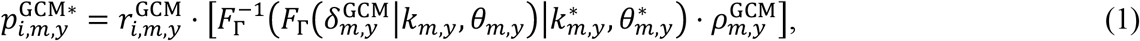

where 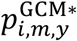 is the bias-corrected precipitation value of the target GCM at year *y*, month *m*, and date *i*. In the equation, 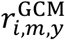 is the ratio of the precipitation value of the GCM relative to the monthly mean value. Then, the following equation is used:

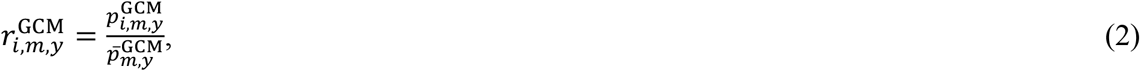

where 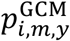 is the precipitation value (not bias-corrected) of the GCM at year *y*, month *m*, and date *i* and 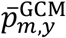 is the monthly mean precipitation value of the GCM at year *y* and month *m*. In equation 1, *F*_Γ_ represents the cumulative distribution function of a gamma distribution, 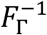 represents the inverse function of the cumulative distribution function of the gamma distribution, and *k*_*m,y*_ and *θ*_*m, y*_ are the shape parameters. In equation 1, 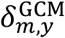 indicates the deviation of the monthly mean from the normal climate value of the corresponding period, and this value calculated as follows:

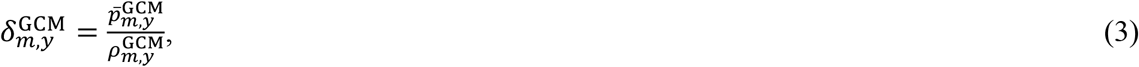

where 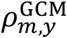 is the normal climate value during the target period. In this method, we defined the normal climate value as the mean of the monthly mean precipitation values over 31 years.

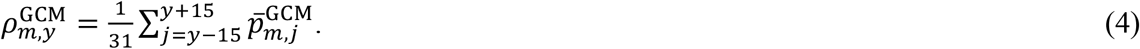

When the mean of a gamma distribution is fixed at one, the shape parameters are represented as follows:

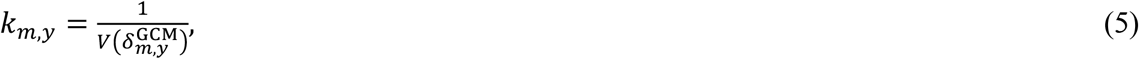

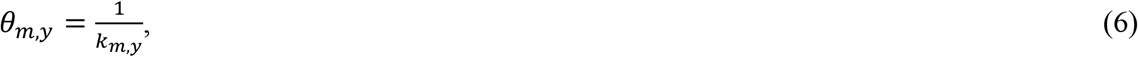

where 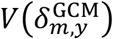 indicates the variance in 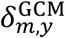 at month *m* over 31 years.

In this method, we assumed that the 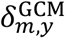 value follows a gamma distribution and that the ratio of the variance of 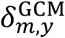 to the variance of 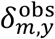 is maintained even in the future scenario. Here, 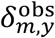 represents the deviation of the monthly mean in the observation data from the normal climate value.

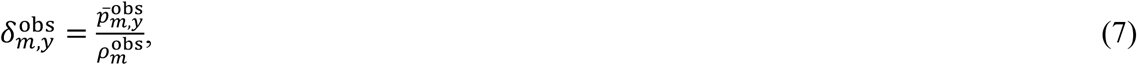

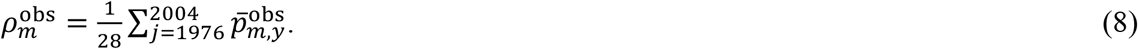

In the above equations, 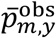 indicates the monthly mean precipitation value in the observed data. As mentioned above, because we assume that the ratio of the variance in 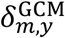 to the variance in 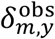 is maintained, 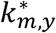 and 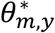 are calculated as follows:

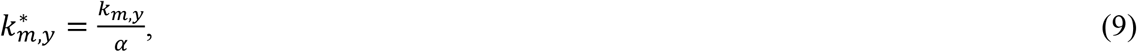

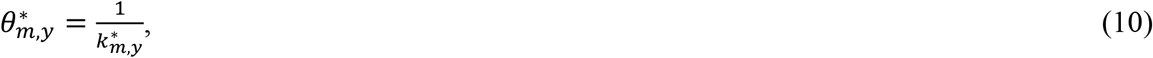

where

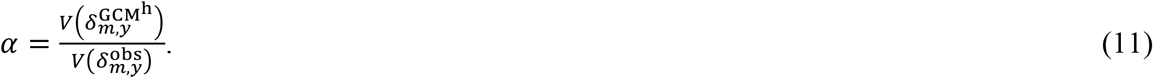

In equation 11, 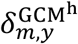 is the deviation of the monthly mean of the historical GCM precipitation data from the normal climate value. Here, we defined the normal climate value as the average monthly mean during 1976–2004.

The method proposed here is an original bias correction method, but the above equations are easily derived if we assume that the 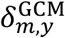 value follows a gamma distribution and that the ratio of the variance in 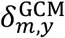 to the variance in 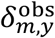 is maintained even in the future scenario. Notably, because we combined this method with the bias correction method described previously^51^, equation 2 should be expressed as follows:

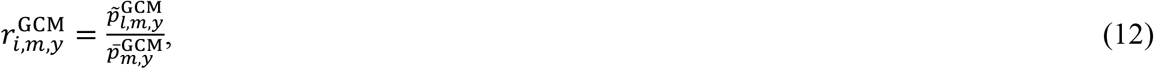

where 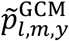 is the precipitation data that are bias-corrected using the method described previously ^51^.

### Analyses

We adopted previously presented^21^ models in which environmental triggers for floral induction accumulate for *n*_1_ days prior to the onset of floral induction (Fig. 3A). Flowers then develop for *n*_2_ days before opening (Extended Data Fig. 2). The model assumption of the time lag between floral induction and anthesis, which is denoted as *n*_*2*_, was validated by a previous finding in which the expression peaks of flowering-time genes, which are used as molecular markers of floral induction, were shown to occur at least one month before anthesis in *Shorea curtisii*^21^. *S. curtissi* is included in our data set. The cool unit at time *t*, CU(*t*|*θ*^*C*^), is calculated as follows:

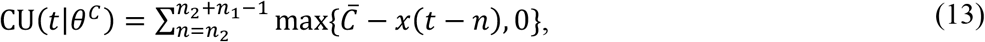

where 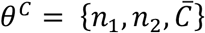 is the set of parameters and *x*(*t*) is the temperature at time *t*. Here, 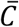 indicates the threshold temperature. The term max{*x*_*1*_, *x*_*2*_} is a function that returns a larger value for the two arguments. Similarly, given 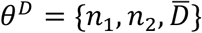, the drought unit at time *t*, DU(*t*|*θ*^*D*^), is defined as the difference between the mean daily accumulation of rainfall over *n*_1_ days and a threshold rainfall level 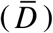:

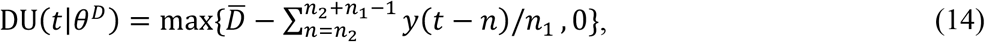

where *y*(*t*) is the rainfall value at time *t*. The term max{*x*_1_, *x*_2_} is defined similarly as in equation 13.

Logistic regression was performed using only the drought unit (DU) and using the product of CU and DU (CU × DU) as the explanatory variables and using the presence or absence of a first flowering event as the dependent variable for each phenological cluster. Because the number of phenological clusters is unknown, we performed forward selection on the cluster number based on the AIC. Let *m* be the number of phenological clusters based on the dendrogram drawn from the time-series clustering explained above (Extended Data Fig. 5). Given *m* phenological clusters, let 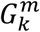 be the *k*th set of clusters in which the DU model is adopted for model fitting. Here, 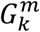 indicates the set of cluster IDs, and *k* ranges from 0 to *m*(*m*+1)/2. For example, when *m* = 2 (i.e., there are two clusters, clusters 1 and 2), there are four cluster sets, calculated as follows:

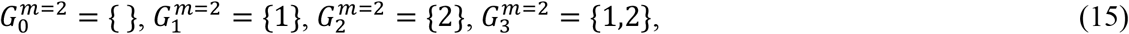

where the element in the bracket indicates the ID of the cluster in which the DU model is adopted for model fitting. When *k* = 0, the DU model is not used; instead, the CU × DU model is adopted for model fitting for both clusters 1 and 2. Let *i* be the *i*th element of the vector **E**, which is defined as follows:

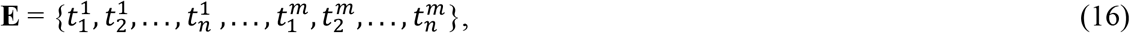

where *n* is the length of the time series data for each cluster. Notably, *n* = 223 is the same for all species and clusters. The term 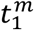 in the above equation denotes the first time point of the time series of length *n* for the species included in cluster *m*. Given *m* and *k*, let *p*^(*m,k*)^(*i*) be the flowering probability of element *i* of vector **E**. The term *p*^(*m,k*)^(*i*) is expressed as follows:

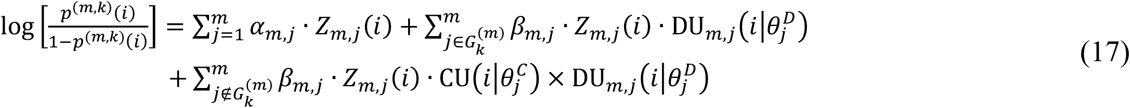

where *Z*_*m,j*_(*i*) is the dummy variable indicating a cluster for *i*; *Z*_*m,j*_ (*i*) equals 1 if the *i*th element of **E** belongs to the *<u>j</u>*th cluster, otherwise it is zero, and *α*_*m,j*_ and *β*_*m,j*_ in equation (5) are regression coefficients for the *j*th cluster when the species are grouped into *m* clusters. We estimate the parameters and the number of clusters based on a finite number of observations. Given the number of clusters *m*, for each of *m* clusters, the parameters were estimated by maximizing the loglikelihood value calculated for all combinations of potential parameter values for *n*_1_, *n*_2_, 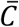, and 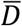 within the ranges of [1 (min), 50 (max)] for *n*_1_, [1, 50] for *n*_2_, [19, 25] for 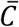, and [1, 9] for 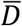. We varied the days (*n*_1_ and *n*_2_) by integers, temperature 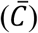 by tenths of a degree C, and daily precipitation 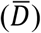 by tenths of a mm. Regression coefficients (*α*_*m,j*_, *β*_*m,j*_) for all *j* values under a given *m* value and associated likelihoods were determined using generalized linear models with binomial error structures.

With the results of the parameter estimations, we determined the number of clusters in two steps. For the first step, for a given *m*, we obtained 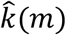 according to the following equation:

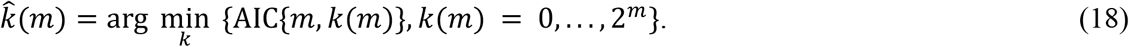

For the second step, with the results of 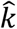 obtained from the first step, we obtained the estimate of the number of clusters according to forward selection by searching for the 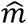 value that satisfies the following inequalities:

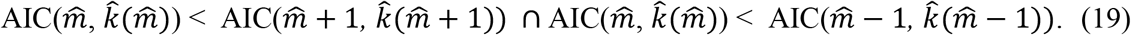

For model fitting, the first flowering month was extracted from the flowering phenology data. When flowering lasted more than one month, the month after the first flowering month was replaced by a value of zero (absence of flowering). If the month before the first flowering month was a missing value, the first flowering month was treated as a missing value and was not used for further analyses. We assumed that phenology monitoring was performed on the first date of each month.

### Projections of 21st century changes in flowering phenology

We used two scenarios (RCP2.6 and RCP8.5) to forecast future reproductive phenology in dipterocarp species for each of the three GCMs (GFDL–ESM2M, IPSL–CM5A-LR, and MIROC5). We predicted the flowering probability per month for each phenological cluster during the periods from May 1, 1976–March 31, 1996 and from January 1, 2050–December 31, 2099 based on the best model (Extended Data Table 3). The predicted flowering probability during the 2050–2099 period was normalized to that during the 1976–1996 period for each climate scenario and for each of three GCMs. To compare the seasonal patterns between 1976–1996 and 2050– 2099, the predicted flowering probability was averaged for each month from January to December and plotted for each month in Fig. 6. R version 3.6.3^52^ was used for all analyses.

## Data availability

The phenology data used in this study will be provided as the Extended Data after acceptance of the paper. The meteorological data provided by the Malaysian Meteorological Department may not be resold, distributed or disclosed to third parties, as regulated by the Malaysian Meteorological Department. Other data that support the findings of this study are available from the corresponding authors (SN and AS) upon reasonable request.

## Code availability

The codes used for model fitting will be provided as Source Data after acceptance of the paper.

## Additional Information

### Supplementary Information

The online version of this paper contains the supplementary materials.

Extended Data 1 | Species list

Extended Data 2 | Order of species listed from the highest to lowest flowering frequency

Extended Data 3 | Results of model fitting

Extended Data 4 | List of dipterocarp species classified into six phenological clusters

**Extended Data Fig. 1.**
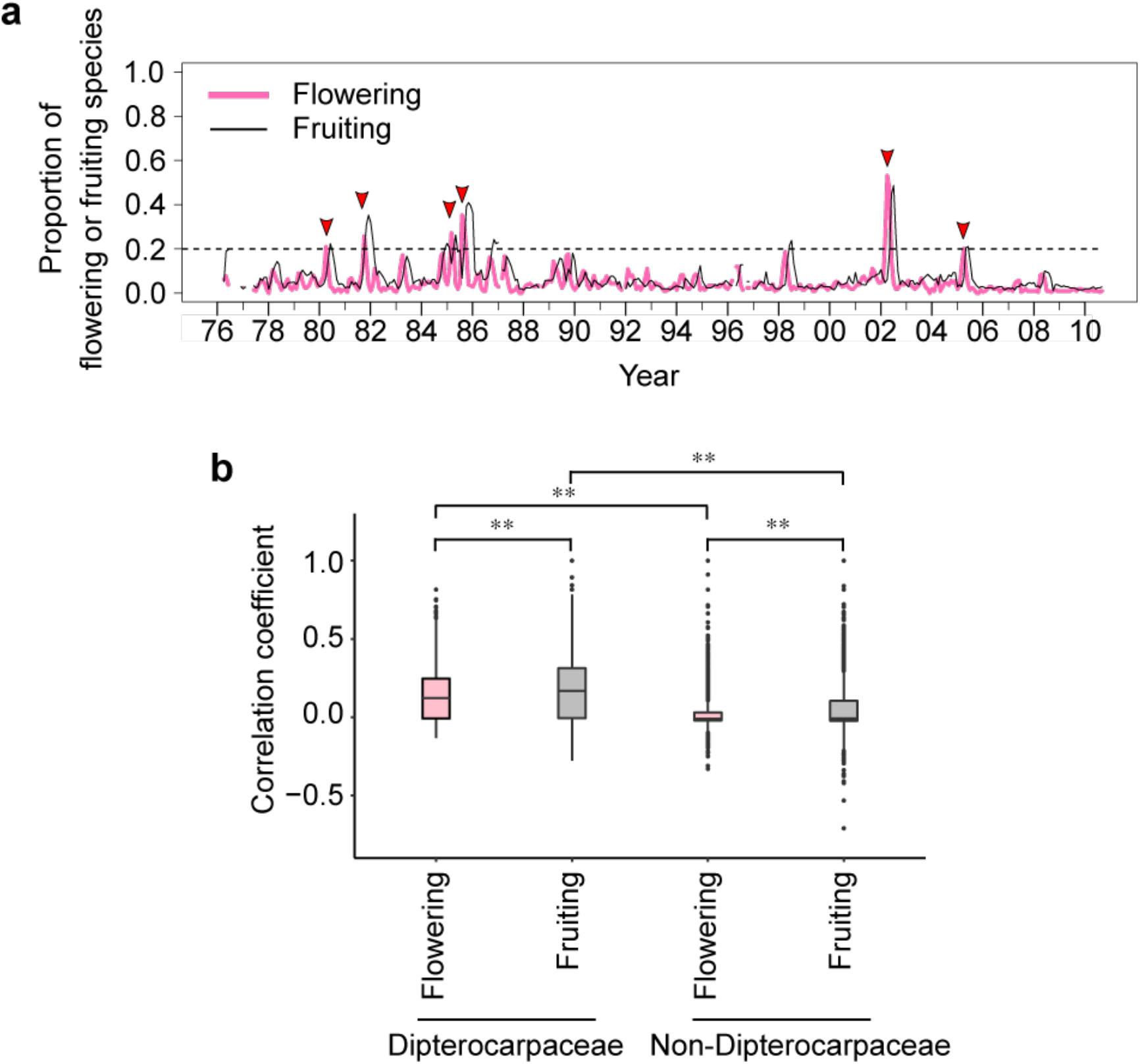
Proportion of flowering and fruiting species in 210 tropical tree species and correlation of flowering and fruiting phenology among species. **a**, The horizontal arrows indicate large flowering events with flowering of more than 20% of the monitored species. A dashed line represents the level of 20% of flowering or fruiting species. **b**, Mean Pearson’s correlation coefficient values of flowering and fruiting binary data over all species pairs, calculated for Dipterocarpaceae (95 species) and non-Dipterocarpaceae species (115 species). Symbols indicate the results of the two-way ANOVA test. ^**^*P* < 0.01. When there were missing values for at least one species, the time point was removed.

**Extended Data Fig. 2.**
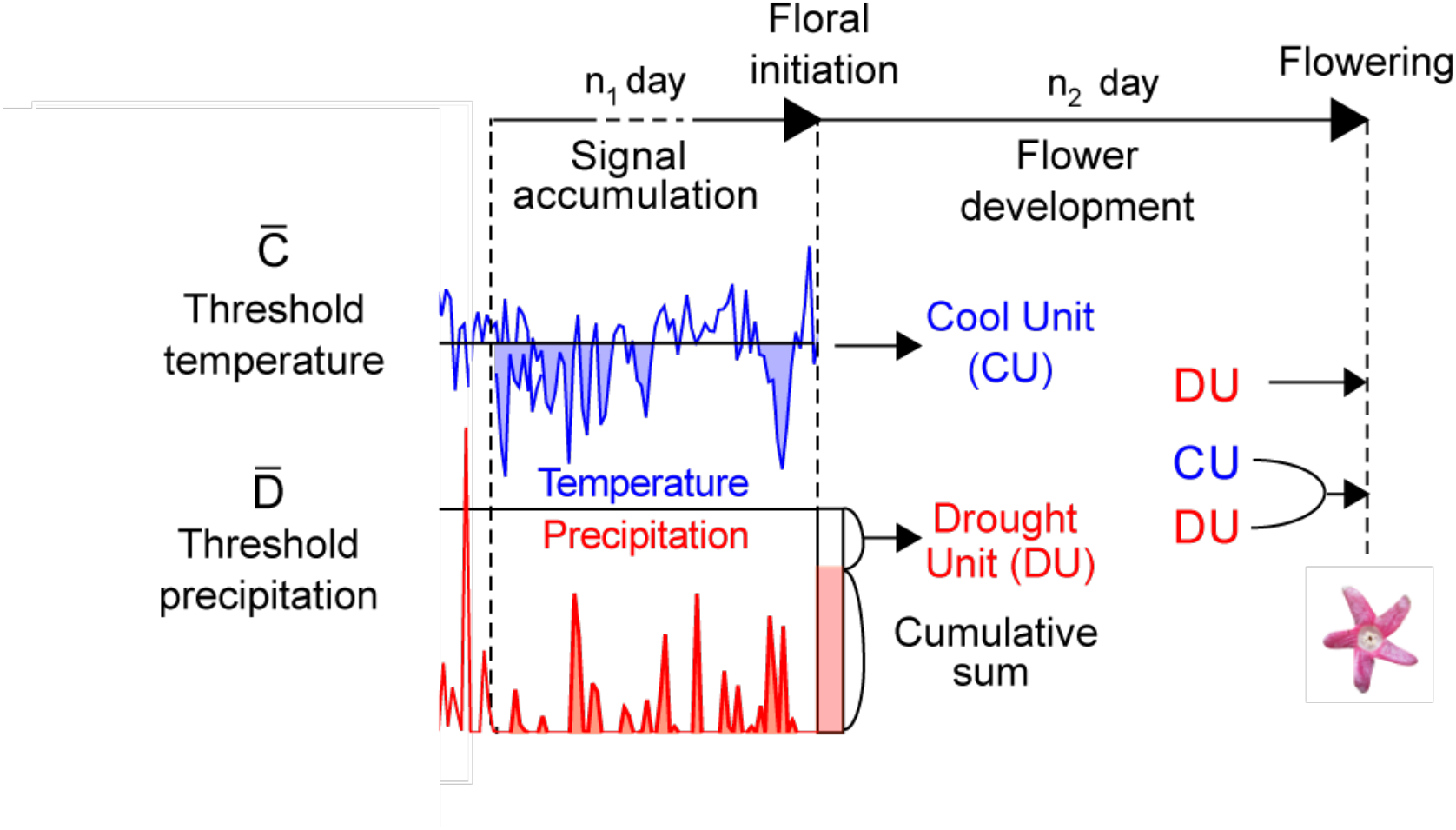
Model structure. The model includes the period of flower development *n*_2_ days after the onset of floral initiation and the accumulation of environmental signals over *n*_1_ days prior to the onset of floral initiation. The cool unit (CU) was equal to the cumulative sum of the negative differences between the threshold temperatures 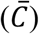 and the daily mean temperatures over *n*_1_ days. The drought unit (DU) equalled the difference between the threshold precipitation level 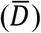 and the cumulative sum of precipitation over *n*_1_ days. Two models were used to describe the relationship between the flowering probability and environmental cues. The two models involved the drought only (DU) and synergistic cold plus drought (CU × DU) models.

**Extended Data Fig. 3.**
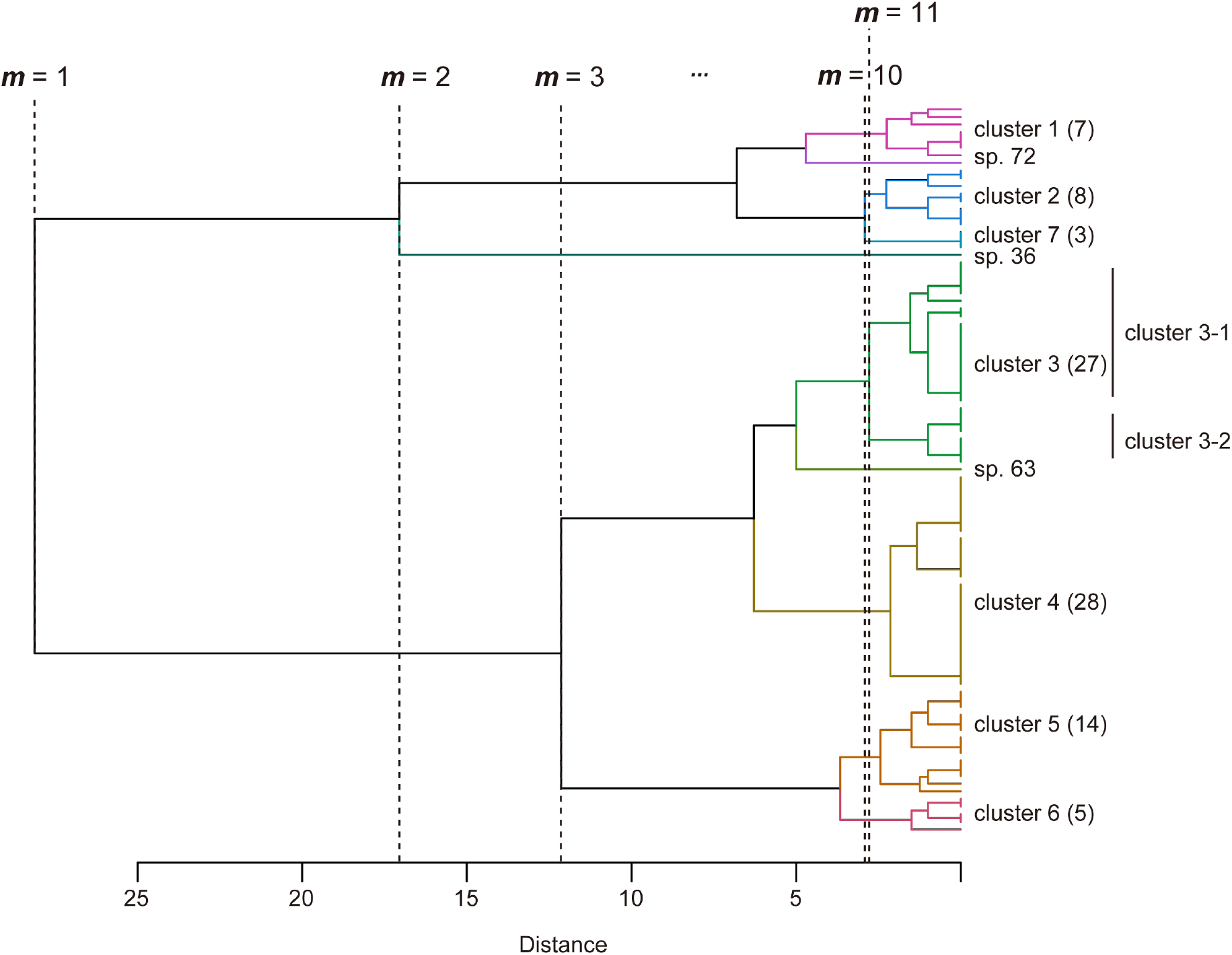
Illustration for the forward selection of the cluster number. A dendrogram indicates the results of the time-series clustering of 95 dipterocarp species. The value *m* is the number of phenological clusters examined for model fitting. The species grouped into the same cluster at *m* = 10 are illustrated by the same colour. The optimal cluster number 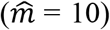 was identified according to the forward selection of optimal 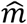 values based on the minimization of the AIC. When *m* = 11, cluster 3 was divided into two subclusters (clusters 3-1 and 3-2). Clusters with fewer than four species (cluster 7 and independent species sp. 36, sp. 63 and sp. 72) were removed before phenology forecasting was conducted due to their small sample size.

**Extended Data Fig. 4.**
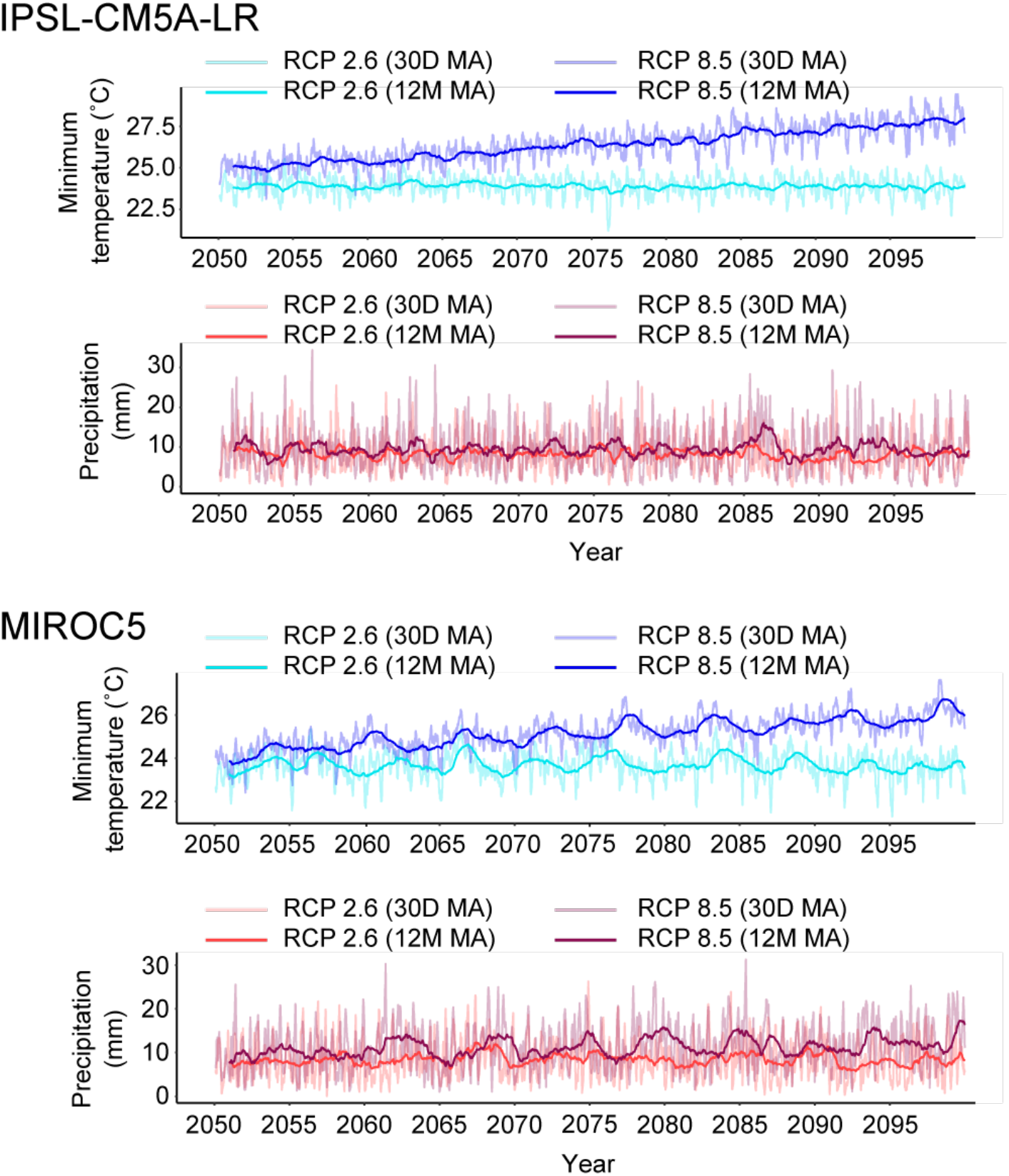
Future minimum temperature and precipitation (2050–2099) values simulated by PSL-CM5A-LR and MIROC5. Future climate conditions were simulated under two climate scenarios (RCP2.6 and RCP8.5). The 30-day (30 D) and 12-month (12 M) moving averages (MAs) were plotted.

**Extended Data Fig. 5.**
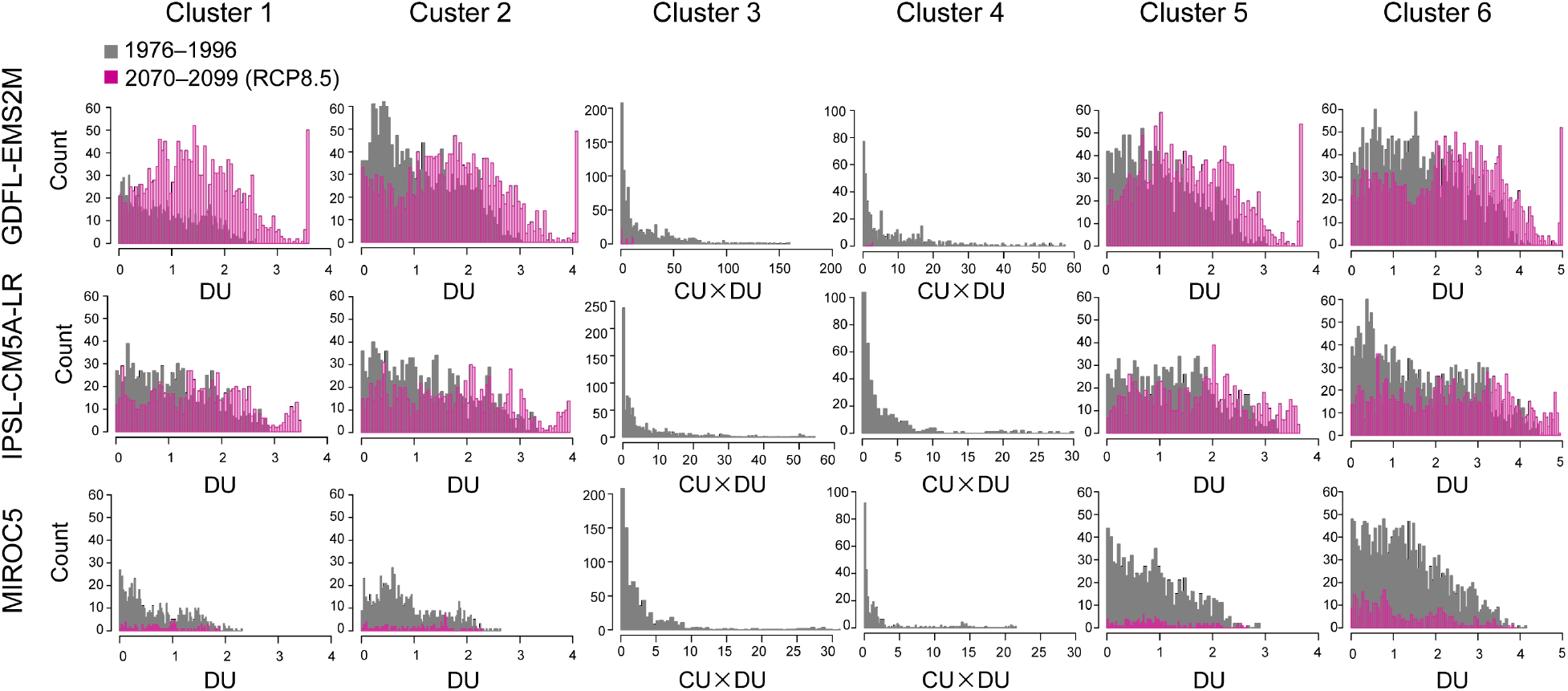
Predicted changes in flowering cues under the RCP8.5 scenario. The predicted environmental signals (predicted with DU or CU × DU) during 1976-1996 (grey) and 2070-2099 (pink) were plotted as histograms for each cluster and for each climate model (GDFL-EMS2M, IPSL-CM5A-LR, and MIROC5).

**Extended Data Fig. 6.**
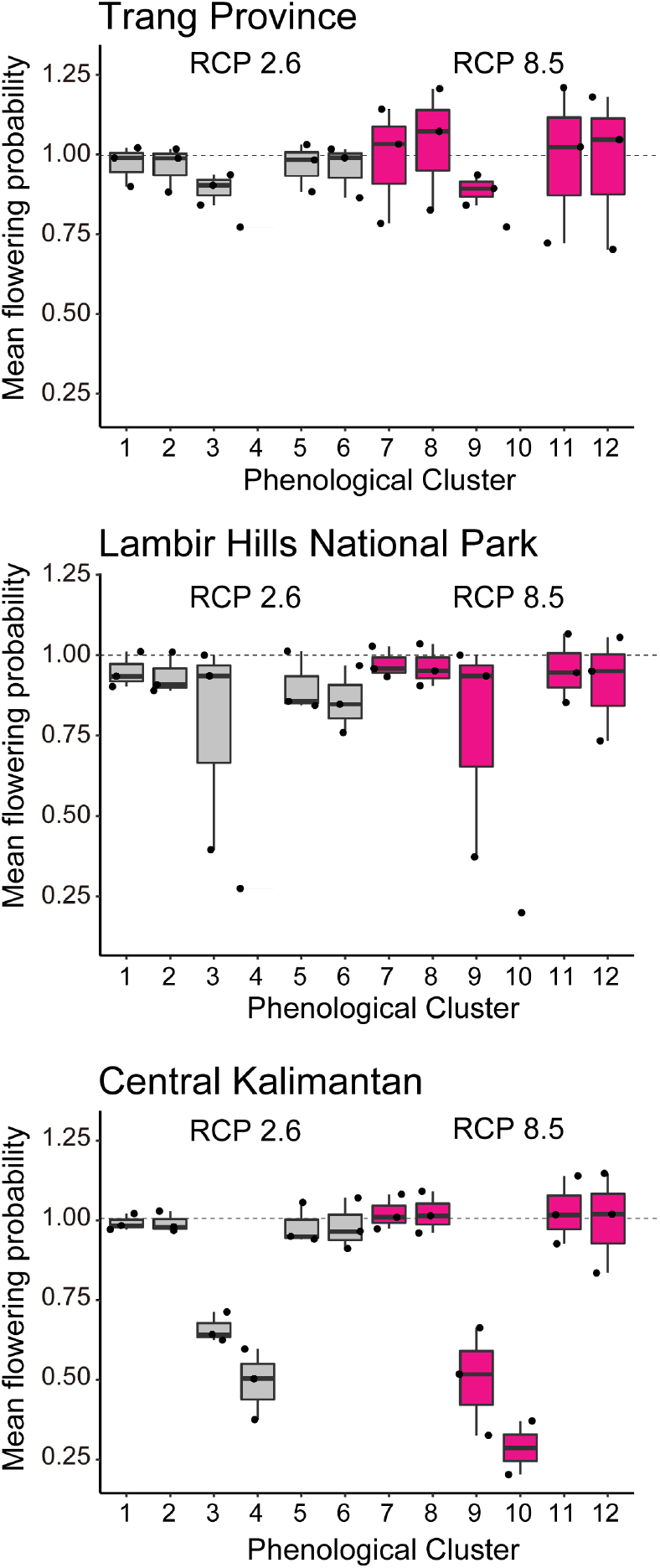
Predictions of future flowering phenology under two climate scenarios (RCP2.6 and RCP8.5) in three regions. The means ± standard errors of the normalized flowering probabilities predicted for 2050–2099 for each phenological cluster under two climate scenarios in three regions (Trang Province, Lambir Hills National Park, and central Kalimantan) are shown. The means and standard errors across three GCMs were calculated from predictions by GDFL–EMS2M, IPSL–CM5A-LR, and MIROC5. For each model and each phenological cluster, the prediction was normalized by the historical climate conditions during 1976–1996. A dotted line indicates the level that is equal to the one for 1976–1996 period. The horizontal line inside each box and the length of the box indicate the median and the interquartile range (the range between the 25th and 75th percentiles), respectively. The whiskers indicate points within 1.5 times the interquartile range. When the maximum flowering probability was less than 0.005 for both periods (1976–1996 and 2050-2099), we omitted the results because of the very low flowering probability.

**Extended Data Fig. 7.**
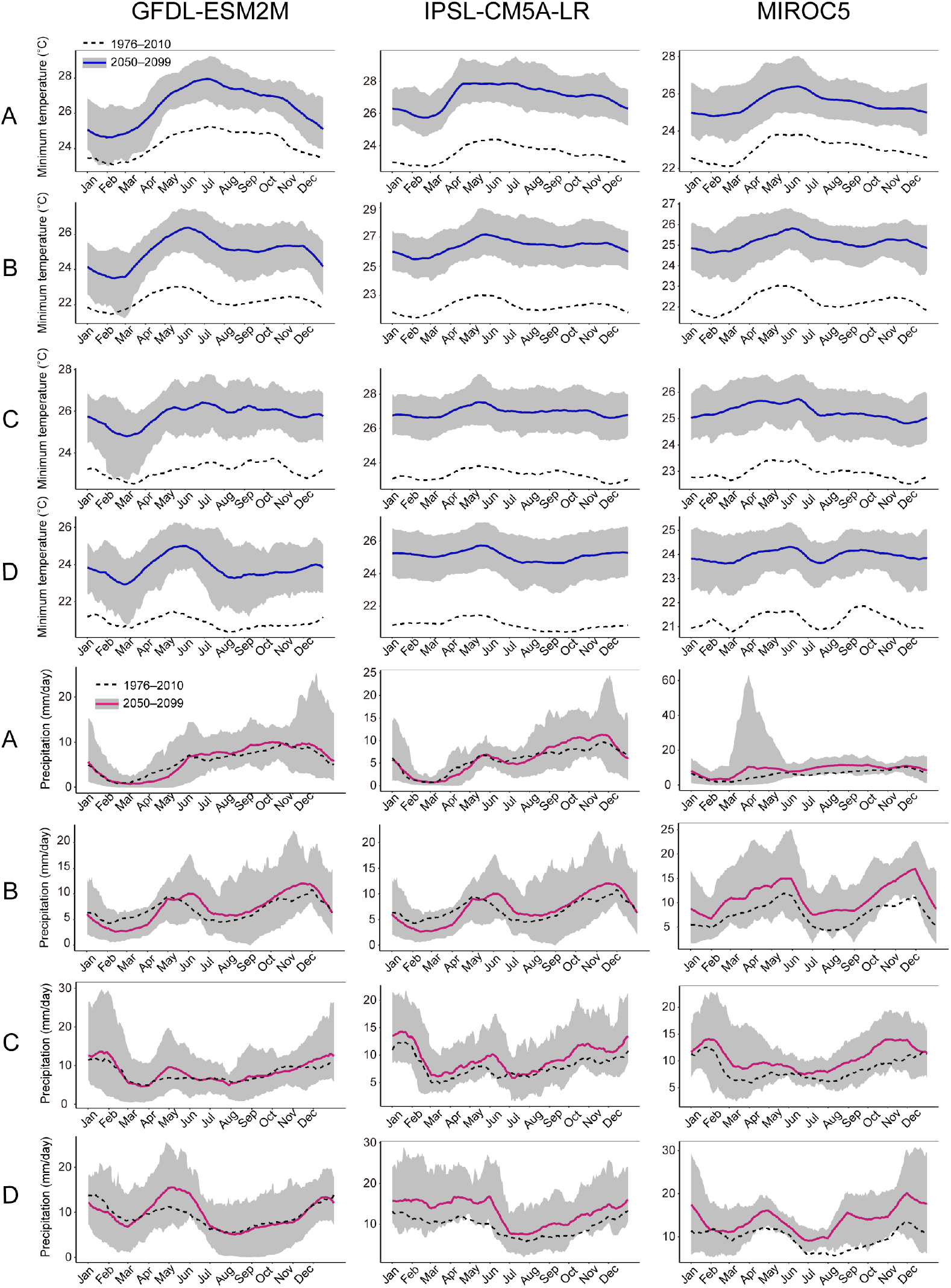
Seasonality in the future climate under the RCP8.5 scenario for each of three GCMs in four regions in Southeast Asia. Averages (blue line) ± standard errors (envelope) of the minimum temperature and averages (pink line) ± standard errors (envelope) of precipitation calculated from the 30-day running means during 2050–2099. Dotted lines indicate the average minimum temperature or precipitation value during the 1976–2010 period. A: Trang province, B: FRIM, C: Lambir Hills National Park, and D: central Kalimantan (see the map in Fig. 6a).

**Extended Data Table 1.** Results of the Mann–Kendall test for detecting long-term trends in the proportions of flowering and fruiting species. The mean change rate was estimated using linear regressions of the monthly flowering or fruiting data at the community level, including 210 tropical tree species.

| Phenological trait | Mann–Kendall $\tau$ | $P$ value | Mean change rate per decade | 2.5% quantile | 97.5% quantile |
| --- | --- | --- | --- | --- | --- |
| Flowering | −0.20 | 2.13E-03 | −0.87E-02 | −1.43E-02 | −3.12E-03 |
| Fruiting | −0.23 | 8.69E-12 | −1.74E-02 | −2.41E-02 | −1.06E-02 |

**Extended Data Table 2.** Parameters and fitting results of the best model. Flowering was assumed to be cued either by the drought unit only (DU) or by a synergism of cool temperatures and drought (CU × DU). In the table, *α* and *β* represent the coefficients of the intercepts and slopes of the logistic regression equations. The best models and parameter values are presented in bold. AUC-train indicates the areas under the ROC curve (AUCs) obtained using training data from June 1976 to March 1996. AUC-valid indicates the AUCs obtained using validation data from July 1997 to April 2005. AUC-valid could not be calculated for cluster 4 because no flowering events were predicted.

| Phenological cluster | Flowering cue | $n_1$<br>(day) | $n_2$<br>(day) | $\bar{D}$<br>(mm) | $\bar{C}$<br>(°C) | $\alpha$ | $\beta$ | Pseudo $R^2$<br>(Nagelkerke) | AUC train | AUC valid |
| --- | --- | --- | --- | --- | --- | --- | --- | --- | --- | --- |
| Cluster 1 | DU | 25 | 38 | 3.6 | – | –2.75 | 1.01 | 0.086 | 0.64 | 0.62 |
| Cluster 2 | DU | 27 | 41 | 4.1 | – | –3.02 | 0.95 | 0.10 | 0.66 | 0.71 |
| Cluster 3 | $CU \times DU$ | 26 | 44 | 4.4 | 20.3 | –4.47 | 0.20 | 0.22 | 0.76 | 0.76 |
| Cluster 4 | $CU \times DU$ | 33 | 19 | 4.8 | 19.4 | –5.93 | 0.66 | 0.18 | 0.70 | n.a. |
| Cluster 5 | DU | 38 | 29 | 3.7 | – | –3.91 | 1.11 | 0.12 | 0.69 | 0.78 |
| Cluster 6 | DU | 38 | 32 | 5.0 | – | –3.86 | 0.93 | 0.20 | 0.78 | 0.79 |

